# Neto proteins differentially modulate the gating properties of *Drosophila* NMJ glutamate receptors

**DOI:** 10.1101/2024.04.22.590603

**Authors:** Tae Hee Han, Rosario Vicidomini, Cathy Isaura Ramos, Mark Mayer, Mihaela Serpe

## Abstract

The formation of functional synapses requires co-assembly of ion channels with their accessory proteins which controls where, when, and how neurotransmitter receptors function. The auxiliary protein Neto modulates the function of kainate-type glutamate receptors in vertebrates as well as at the *Drosophila* neuromuscular junction (NMJ), a glutamatergic synapse widely used for genetic studies on synapse development. We previously reported that Neto is essential for the synaptic recruitment and function of glutamate receptors. Here, using outside-out patch-clamp recordings and fast ligand application, we examine for the first time the biophysical properties of recombinant *Drosophila* NMJ receptors expressed in HEK293T cells and compare them with native receptor complexes of genetically controlled composition. The two Neto isoforms, Neto-α and Neto-β, differentially modulate the gating properties of NMJ receptors. Surprisingly, we found that deactivation is extremely fast and that the decay of synaptic currents resembles the rate of iGluR desensitization. The functional analyses of recombinant iGluRs that we report here should greatly facilitate the interpretation of compound *in vivo* phenotypes of mutant animals.

## Introduction

Ionotropic glutamate receptors (iGluRs) mediate fast excitatory synaptic signaling throughout the vertebrate CNS and also at the neuromuscular junction (NMJ) of insects and crustaceans (Takeuchi and Takeuchi 1964; Jan and Jan 1976; Traynelis et al. 2010; Yu et al. 2021). iGluRs are tetrameric channels that achieve strikingly diverse biophysical properties by combining different iGluR subunits within a receptor complex and by association with a rich array of auxiliary subunits (Jackson and Nicoll 2011; Hansen et al. 2021). Auxiliary subunits bind to iGluRs at many stages of the receptor life-cycle and modulate not only channel properties but also the delivery of receptors to the cell surface, their subcellular distribution, synaptic recruitment, and association with various postsynaptic density (PSD) scaffolds. In *Drosophila*, multiple NMJ iGluR subunits (DiAntonio 2006), and the auxiliary protein Neto (Neuropillin and Tolloid-like) are each essential for viability (Kim et al. 2012), indicating that Neto is required for function of the NMJ. Phylogenetic studies indicate that fly NMJ iGluRs belong to the kainate receptor (KAR) clade, which has been expanded in *Diptera* (Li et al. 2016). Since Neto appears to be a KAR-dedicated auxiliary subunit (Zhang et al. 2009; Tomita and Castillo 2012), it would be anticipated that similar to vertebrate KARs, *Drosophila* Neto modulates the gating of NMJ iGluRs. However, the biophysical properties of native *Drosophila* iGluRs and their modulation by auxiliary proteins remain poorly understood. In flies as in humans, synapse strength and plasticity is determined by the interplay between expression of different iGluR subtypes (DiAntonio et al. 1999). At the fly NMJ, type-A and type-B iGluRs are assembled from four different subunits: either GluRIIA (type-A) or GluRIIB (type-B), plus one copy each of GluRIIC, GluRIID and GluRIIE (Petersen et al. 1997; DiAntonio et al. 1999; Marrus et al. 2004; Featherstone et al. 2005; Qin et al. 2005). The shared subunits and Neto are both essential for viability and for iGluR synaptic recruitment (DiAntonio 2006; Kim et al. 2012). In flies, Neto and iGluRs depend on each other for trafficking and stabilization at synaptic sites (Kim et al. 2012). Neto proteins are single-pass transmembrane proteins with a highly conserved extracellular part containing two C1r/C1s–Uegf–BMP domains (known as CUB1 and CUB2) and a low-density lipoprotein class A motif (LDLa) and variable intracellular domains. Recent cryo-electron microscopy studies indicate that the vertebrate Neto2 accesses different interfaces of GluK2 homotetrameric receptors, making tight inter-subunit connections via CUB1/GluK2-ATD and LBD domains and intra-subunit interactions within the transmembrane helices (He et al. 2021). In addition to the conserved protein interaction domains, *Drosophila* Neto has multiple intracellular putative docking motifs, and phosphorylation sites indicating that Neto likely controls NMJ iGluR synaptic recruitment and function via binding to iGluRs, to PSD components, as well as other interacting partners (Kim et al. 2015; Ramos et al. 2015; Sulkowski et al. 2016; Han et al. 2020). The *Drosophila neto* gene codes for two isoforms (Neto-α and Neto-β) with completely different intracellular domains of 206 and 351 residues, generated by alternative splicing (Ramos et al. 2015). Either isoform can sustain organism viability, but they have different *in vivo* distributions and functional roles (Kim et al. 2012; Han et al. 2015; Kim et al. 2015; Ramos et al. 2015; Han et al. 2020).

We previously reported the functional reconstitution of recombinant *Drosophila* NMJ iGluRs in *Xenopus* oocytes (Han et al. 2015). These studies showed that just as in flies, four different subunits, GluRIIA or GluRIIB, plus GluRIIC, GluRIID and GluRIIE are required for the robust surface expression of *Drosophila* NMJ iGluRs; complexes assembled from fewer than four subtypes do not reach the cell surface, and remain trapped in secretory compartments. Our experiments also revealed that in the absence of Neto *Drosophila* NMJ iGluRs are not activated by glutamate even after application of the lectin Concanavalin A (Han et al. 2015), thus establishing that with heterologous expression we can recapitulate results obtained *in vivo* using fly genetics. However, in contrast to vertebrate iGluRs, for which the kinetics of activation, deactivation and desensitization have been studied in depth, with extensive characterization of the properties of different subunit combinations and auxiliary proteins (Hansen et al. 2021), comparable studies on recombinant *Drosophila* iGluRs have not been reported. This is a big gap in the field of developmental neurobiology: the *Drosophila* NMJ has been a powerful system to study glutamatergic synapse assembly, development and homeostasis and extensive genetic manipulations of wild-type and mutant NMJ iGluR subunits contributed to our understanding of underlying molecular mechanisms. For example, studies of flies with *GluRIIA* mutations at sites that alter the kinetics of desensitization of mammalian GluA2 AMPA receptors revealed impaired trafficking behavior *in vivo* and abnormal distribution at postsynaptic densities (PSDs) (Petzoldt et al. 2014). However, because the recombinant expression of *Drosophila* NMJ GluRs had not been established, the gating behavior of these *Drosophila* NMJ iGluR variants was not determined, and was assumed to mimic that of their mammalian counterparts.

Here we examine the gating properties of *Drosophila* NMJ iGluRs using outside out patch recordings from HEK293T cells transfected with different combinations of NMJ iGluR subunits and Neto splice variants. In addition, we generated flies with genetically controlled subunit composition and recorded responses from outside out patches obtained from *Drosophila* larval muscle, as well as excitatory synaptic currents from their NMJ. We used rapid application of glutamate to examine the kinetics of activation, deactivation and desensitization, the effect of polyamine toxins, and the effect of the lectin Concanavalin A. We find that the two Neto isoforms differentially modulate the deactivation and desensitization of type-A and type-B receptors. Our study reveals that *Drosophila* Neto is not only required for channel function but also increases the repertoire of channel properties.

## Results

### *Drosophila* NMJ iGluRs form Neto-dependent rapidly desensitizing receptors

To facilitate cell surface expression of recombinant *Drosophila* NMJ iGluRs, we replaced endogenous signal peptides with an optimized sequence and added a C-terminal RGSH6 epitope to all iGluR constructs. Four receptor subunits (GluRIIC, GluRIID, GluRIIE and either GluRIIA or GluRIIB) were transiently transfected in HEK293T cells with or without Neto splice variants and incubated at 30°C for three days prior to outside-out patch recordings. Since these receptors have low sensitivity to glutamate (Heckmann et al. 1996; Han et al. 2015) we examined their gating properties in response to rapid application of 10 mM glutamate. In the absence of Neto, we observed no response to glutamate in 79 patches of GluRIIA/C/D/E and 46 patches of GluRIIB/C/D/E (Figure 1A-B). However, when type-A receptor complexes were co-transfected with *Drosophila* Neto variants (Figure 1C), a large fraction of outside-out patches (78/102) yielded macroscopic currents in response to glutamate. A similar requirement for Neto was observed for type-B receptors. Rapid application of 10 mM glutamate to outside-out patches from HEK cells transfected with various iGluR/Neto combinations revealed subunit composition-dependent gating properties and differential effects of Neto variants (Table 1). For example, for macroscopic currents recorded from type-A iGluR/Neto-α (A/α) complexes the 10%-90% rise time was 330 ± 9.64 µs (n = 8), with a deactivation time constant (τ_off_) 0.64 ± 0.07 ms, whereas type-A iGluR/Neto-β (A/β) complexes had a similar rise time, 340 ± 17.77 µs (n = 8), but 1.4 fold slower deactivation, τ_off_ 0.90 ± 0.11 ms (Figure 1D). Longer applications of glutamate (100 ms) revealed rapid and profound desensitization of more than 98% for both complexes (n = 7-8), with the decay best fit by the sum of two exponential functions: A/α τ_fast_ 2.01 ± 0.28 ms, τ_slow_ 5.92 ± 1.07 ms, A_fast_ 81.49 ± 4.61 %; and A/β τ_fast_ 3.83 ± 0.84 ms, τ_slow_ 10.37 ± 2.0 ms, A_fast_ 80.02 ± 6.84 % (Figure 1E).

**Figure 1.**
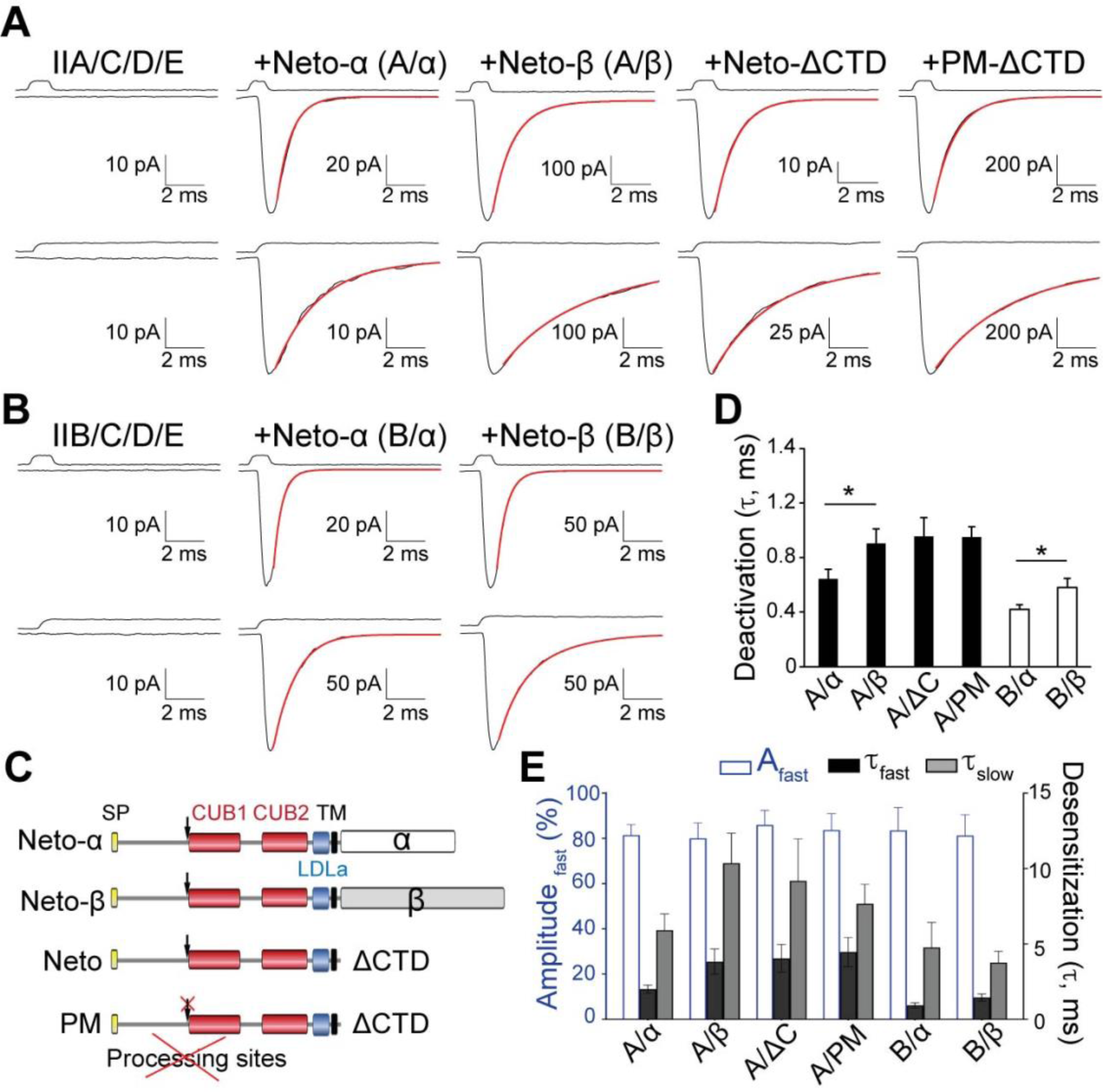
Differential modulation of *Drosophila* NMJ iGluRs by Neto isoforms. (**A-B**) Responses to 10 mM glutamate applied for 1 ms (upper traces) and 100 ms (lower traces) to outside-out patches from HEK293T cells transfected with GluRIIA/C/D/E (**A**) and GluRIIB/C/D/E (**B**) without Neto or in the presence of different Neto splice variants. Black lines show the average of 35-60 responses from one patch; red lines show the decay of the responses fitted with the sum of one (upper) or two (lower) exponential functions; open tip junction currents measured at the end of the experiments are shown at the top. The holding potential was - 60mV for all recordings. (**C**) Diagram of Neto variants utilized. *Drosophila* Neto isoforms are expressed as pre-proteins; their inhibitory pro-domain must be cleaved at conserved Furin processing sites (marked by arrows) before Neto can promote formation of synaptic iGluR aggregates in vivo. PM denotes a processing mutant unable to shed the inhibitory pro-domain. (**D-E**) Summary graphs for deactivation, *p<0.05, (**D**) and desensitization time constants (**E**) for various iGluR/Neto complexes; the A_fast_ (%) component for desensitization is shown in blue.

**Table 1.**
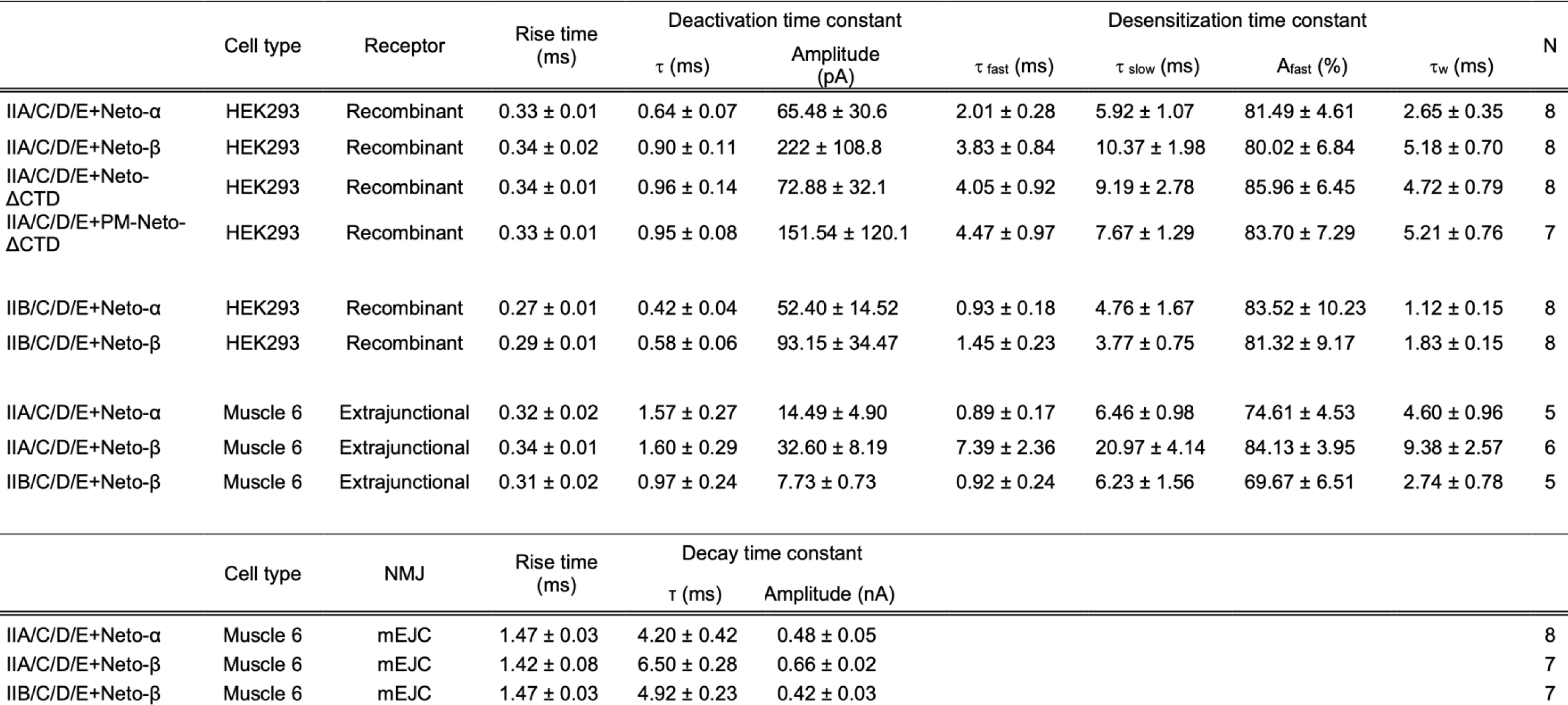
Upper segment: Kinetic analysis for macroscopic currents evoked by 1 ms (deactivation) and 100 ms (desensitization) applications of glutamate recorded using either outside out patches from HEK cells transfected with the indicated iGluR subunit combinations and Neto splice variants (Recombinant) or patches obtained from muscle 6 for larvae with a defined subunit composition (Extrajunctional). Lower segment: Kinetic analysis for mEJCs recorded using 2-electrode voltage clamp for larvae with a defined subunit composition.

Differential modulation by Neto isoforms was also observed for type-B receptors (Figure 1B): These receptors had 10%-90% rise times of 267 ± 9 μs and 294 ± 10 μs for B/α and B/β respectively, with deactivation time constants of 0.42 ± 0.04 ms and 0.58 ± 0.07 ms (Figure 1D). Similar to type-A iGluRs, type-B receptors in complexes with Neto-α also desensitized faster than for Neto-β, with B/α τ_fast_ 0.93 ± 0.18 ms, τ_slow_ 4.76 ± 1.67 ms, A_fast_ 83.52 ± 10.23 %; and B/β τ_fast_ 1.45 ± 0.23 ms, τ_slow_ 3.77 ± 0.75 ms, A_fast_ 81.32 ± 9.17 % (Figure 1E). Of note, the deactivation and desensitization time constants for type-A iGluR/Neto-ΔCTD complexes, τ_off_ 0.96 ± 0.14 ms; desensitization τ_fast_ 4.05 ± 0.92 ms, τ_slow_ 9.19 ± 2.78 ms, A_fast_ 85.96 ± 6.45% were comparable to type-A iGluR/Neto-β complexes but different from type-A iGluR/Neto-α (Figure 1C-E). This indicates that a “minimal Neto”, which retains the highly conserved extracellular and transmembrane domains, but lacks any intracellular parts, is sufficient for NMJ iGluR function. These results also reveal that the cytoplasmic domain of Neto-α, but not of Neto-β, modulates the gating properties of NMJ iGluR receptors, increasing the rate of deactivation and desensitization. Similar results were obtained for type-A receptors in complex with PM-Neto-ΔCTD (Figure 1C-E), a processing mutant variant that cannot shed its inhibitory pro-domain and fails to cluster iGluRs *in vivo* (Kim et al. 2015). This indicates that responses observed in outside-out patches are likely independent of receptor clustering. To allow comparison with prior studies on native extrajunctional *Drosophila* NMJ iGluRs, for which desensitization time constants with single exponential fits of 17.5, 18.8 and 2.0 ms were reported for wild type, type-A and type-B, respectively (DiAntonio et al. 1999), a weighted value (τ_w_) was calculated (see Methods). We find that, type-A recombinant NMJ iGluRs channels have faster desensitization compared to native wild type and native type-A receptors, with weighted decay time constants τ_w_ A/α, 2.65 ± 0.35 ms and A/β, 5.18 ± 0.70 ms, while the rate of desensitization of B/α, 1.12 ± 0.15 ms and B/β, 1.83 ± 0.15 ms, respectively, is comparable to native type-B iGluRs at the NMJ.

In many recordings from patches with macroscopic currents we observed trial to trial fluctuations in amplitude for both short (1 ms) and long (100 ms) applications of glutamate. Given the large number of channels in each patch (around 10-50), these fluctuations likely occur because the open probability (Po) is relatively low. We examined this possibility using non stationary analysis of variance (Supplemental figure S1A-D, quantified in S1E-F). In many patches the open probability was too low to accurately estimate, range (0.1-0.6), consistent with a prior study that reported a value of 0.4 estimated from binomial analysis of single channel current amplitudes (Heckmann et al. 1996). Estimates of the single channel conductance obtained from non-stationary analysis of variance, 160 ± 23 pS, 164 ± 22 pS and 149 ± 39 pS for A/α, A/β and B/α, respectively, gave values comparable to those reported previously from single channel recordings from extra-junctional NMJ iGluRs (Chang et al. 1994; DiAntonio et al. 1999). In many patches with macroscopic currents we observed steps corresponding to the closure or desensitization of individual channels (Supplemental figure S1A-D).

**Supplemental figure S1.**
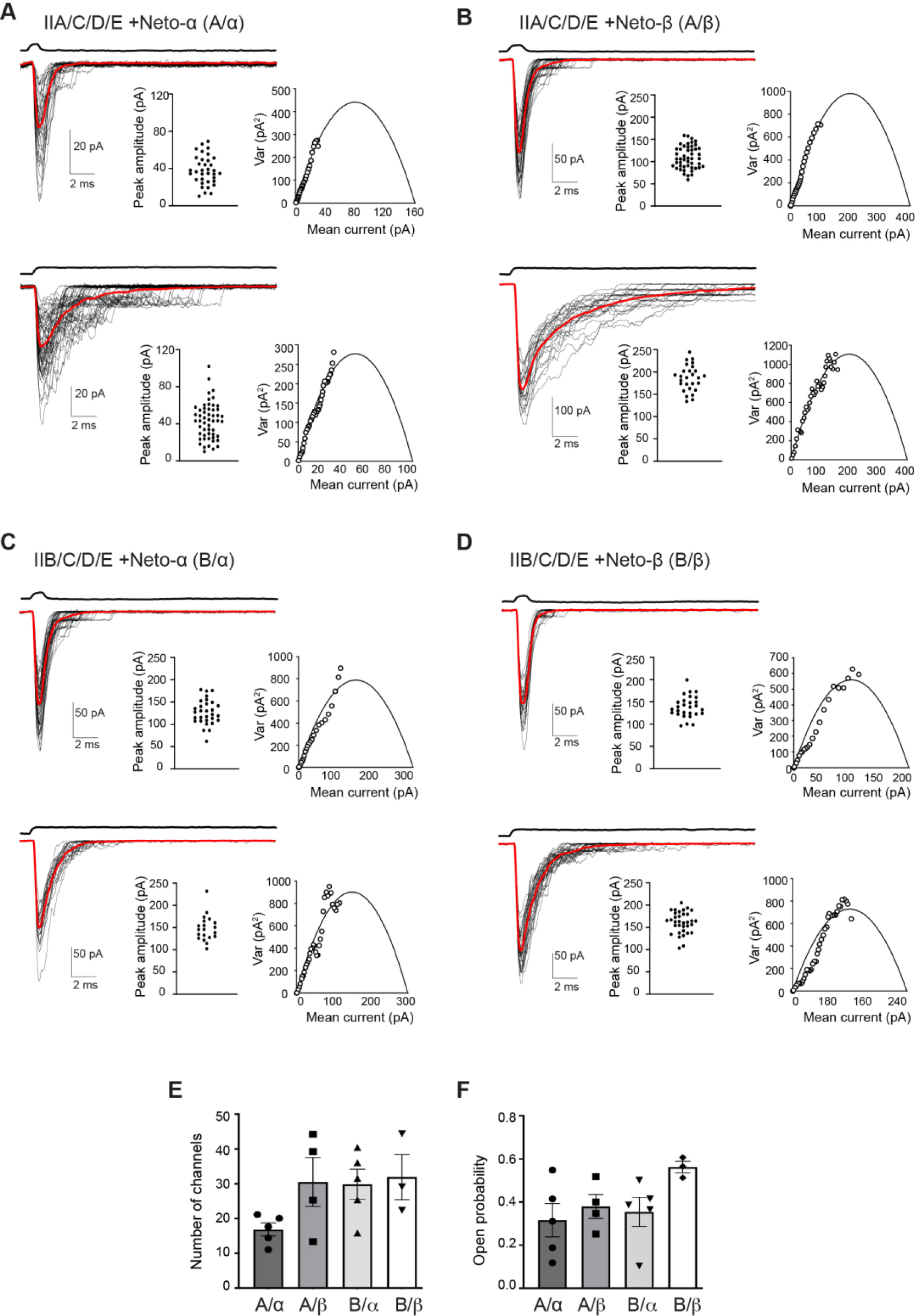
Nonstationary analysis of variance of responses to 10 mM glutamate applied for 1 ms (upper traces) and 100 ms (lower traces) to outside-out patches from HEK cells transfected with various iGluR/Neto receptor complexes as indicated. (**A-D**) Superimposed individual responses are shown in black; the average current in red; open tip junction currents measured at the end of the experiments are shown at the top. The holding potential was - 60mV for all recordings. The insets in each panel show the peak amplitude for all trials (left) and the current-variance relationship (right) fit with the function σ^2^=*i*I - I^2^/N, where σ^2^ is the variance, i is the mean current, N is the number of channels. (**E**) The mean number of channels and the open probability (**F**) determined by variance analyses of responses for 3-5 patches for different iGluR/Neto channel complexes as indicated, showing responses for individual patches and the mean ± SEM.

### Modulation of *Drosophila* NMJ iGluR single channel responses by Concanavalin A

The lectin Concanavalin A (Con A) attenuates iGluR desensitization for a wide variety of species, including locust NMJ iGluRs (Mathers and Usherwood 1976), native vertebrate kainate receptors (Huettner 1990; Wong and Mayer 1993) and AvGluR1 from the primitive eukaryote *Adineta vaga* (Lomash et al. 2013). In prior work, which used *Xenopus* oocytes to study recombinant *Drosophila* NMJ iGluRs, whole cell responses to glutamate were only detectable after treatment with Con A. In the present study used single channel recording to study the effects of Con A. We found that by reducing the incubation time after transfection from three to two days, we could obtain patches where only single channel currents could be detected (Figure 2A-B). For control patches, the single channel conductance at -60 mV measured using individual openings was A/α 175 ± 12 pS (n=14), A/β 172 ± 11 pS (n=16), B/α 173 ± 12 pS (n=10) and B/β 169 ± 9 pS (n=11), similar to estimates from nonstationary analysis of variance (Figure 1-figure supplement 1). From averages of 16-57 responses from single channel patches we estimated desensitization time constants by fitting single exponentials: τ_des_ A/α 2.04 ± 0.32 ms (n=6), τ_des_ A/β 3.71 ± 0.40 ms (n=8), τ_des_ B/α 1.05 ms (n=2) and τ_des_ B/β 1.54 ± 0.20 ms (n=3); these values are comparable to those obtained for macroscopic currents recorded from multichannel patches (Figure 1 and Table 1) with faster desensitization for type-B *versus* type-A and for Neto-α *versus* Neto-β.

**Figure 2.**
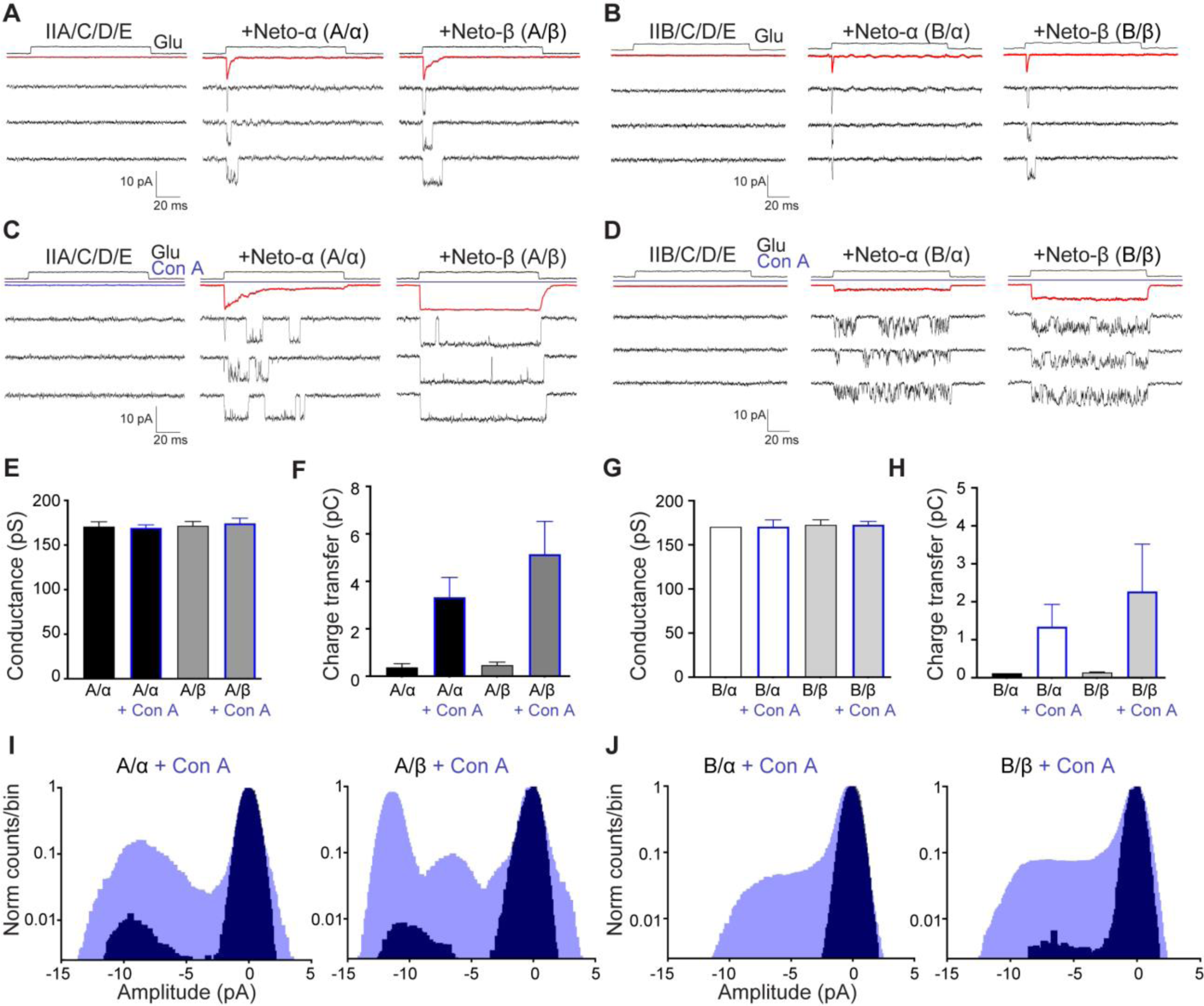
Concanavalin A attenuates desensitization and increases single channel activity evoked by glutamate. (**A** and **C**) Responses for outside-out patches to 10 mM glutamate applied for 100 ms to HEK293T cells transfected with type-A receptors with or without Neto splice variants, as indicated, before (**A**) and after (**C**) treatment with 0.6 mg/ml Con A for 10 min (blue line); average responses (top, red line) and three single channel responses from one patch are shown for each channel variant. The same sequence is shown for type-B receptors (**B** and **D**). The holding potential was -60 mV for all recordings. Open tip junction currents measured at the end of the experiments are shown at the top. (**E-F**) Single channel conductance at -60 mV and the charge transfer for responses to glutamate measured by integration of 35-55 trials shown in panels (**A** and **C**) for type-A receptors. (**G-H**) show the same analysis for type-B receptors. (**I-J**) All points amplitude histograms plotted on a log scale before (black) and after (blue) treatment with Con A, normalized to the closed state peak at 0 pA. Note the shift in the peak profiles at -10 pA (open channels) after application of Con A and the asymmetric profile for the open state. Data are represented as mean ± SEM.

To study the effect of Con A on single channel activity, we treated HEK cells transfected with iGluR/Neto combinations with 0.6 mg/ml Con A for 10 min, and then excised outside-out patches to record responses to 10 mM glutamate applied for 100 ms. We found that patches from HEK cells treated with Con A were less stable, and that it was more difficult to obtain giga-ohm seals, but in exceptional cases sufficient data was obtained for kinetic analysis. In control patches not treated with Con A, single channel events were observed only at the start of the application of glutamate (Figure 2A-B) reflecting the rapid onset of desensitization observed for macroscopic responses (Figure 1). By contrast, pretreatment with Con A dramatically increased single channel activity, revealing major differences between type-A and type-B receptors and also between Neto splice variants. For A/α the response to glutamate showed much slower onset desensitization, time constant 15.5 ms (range 10-26 ms, n=3) 6-fold slower after treatment with Con A *versus* the value for control patches, time constant 2.65 ms, with very few openings 70 ms after the start of the application of glutamate (Figure 2C). By contrast, for A/β, B/α, and B/β there was a complete block for some patches, while the average current revealed variable extents of desensitization from patch to patch (Figure 2C-D). After treatment with Con A single channel activity for A/α and especially A/β consisted of bursts of long duration openings, with few closures within a burst (Figure 2C); by contrast, bursts for B/α and B/β were interrupted by brief closures at high frequency (Figure 2D). Following termination of the application of glutamate, the average response after treatment with Con A showed tail currents with fast decays, time constant A/α 1.9 ms ± 0.2 ms (n=4); A/β 3.8 ms ± 0.4 ms (n=5); B/α 0.9 ms ± 0.2 ms (n=4); B/β 1.2 ms ± 0.1 ms (n=3), suggesting that even after application of Con A glutamate dissociates rapidly following channel closure.

The effects of Con A occurred without any change in single channel conductance (Figures 2E and 2G). Calculation of the charge transfer by integration of raw data in response to application of glutamate for 100 ms revealed substantial differences between type-A and type-B receptors, and also between Neto-α *versus* Neto-β. For control patches the charge transfer was A/α 0.37 ± 0.16 pC (n=4), A/β 0.47 ± 0.12 pC (n=8), B/α 0.12 pC (n=2) and B/β 0.14 ± 0.01 (n=3) pC, with a large increase for patches treated with Con A (Figures 2F and 2H), charge transfer A/α 3.33 ± 0.83 pC (n=5), A/β 5.14 ± 1.38 pC (n=6), B/α 1.34 ± 0.59 pC (n=5) and B/β 2.27 ± 1.25 pC (n=5).

Estimates of open probability were obtained from analysis of all point amplitude histograms for pooled data from 2-8 patches (Figure 2I-J). Because the open state histograms were not well fit by single gaussian functions, we calculated the open probability (Po) by integration which gave control values of 0.039, 0.044, 0.015 and 0.016 for A/α, A/β, B/α and B/β and 0.37, 0.55, 0.20 and 0.29 for patches treated with Con A (Figure 2I-J).

### Single channel kinetics and subconductance states

Closer inspection of single channel records for A/α and A/β complexes revealed bursts of openings, during which the current fluctuated between well-defined subconductance states with amplitudes of 50% and 75% of the open state conductance (Figure 3A-B); we did not observe openings to 25%, but it is possible these might occur with lower concentrations of glutamate. Using a cutoff time of 200 µs, open time and burst duration histograms for A/α and A/β were well fit by the sum of two exponentials for both control patches, and for patches from HEK293T cells pretreated with Con A (Figures 3C, 3E, and Supplemental Table 1). For A/α this analysis revealed a 2-4 fold increase in the duration of the slow component of the open time, 3.75 *versus* 7.53 ms, and burst length, 3.12 *versus* 11.53 ms, for control and Con A respectively. For A/β the duration of the fast component of the burst length, 1.28 *versus* 3.32 ms, and open time, 1.11 *versus* 5.60 ms, also increased 2.6 and 5-fold, respectively for Con A treated patches (Figure 3D). Notably, for type-A/β the duration of the slow component of the open time and burst length distribution increased even more dramatically, from control values of 4.51 and 4.38 ms, to events of duration longer than 100 ms (Figure 3E and Supplemental Table 1), with for many trials channel closure only after the termination of the application of glutamate (Figure 2C).

**Figure 3.**
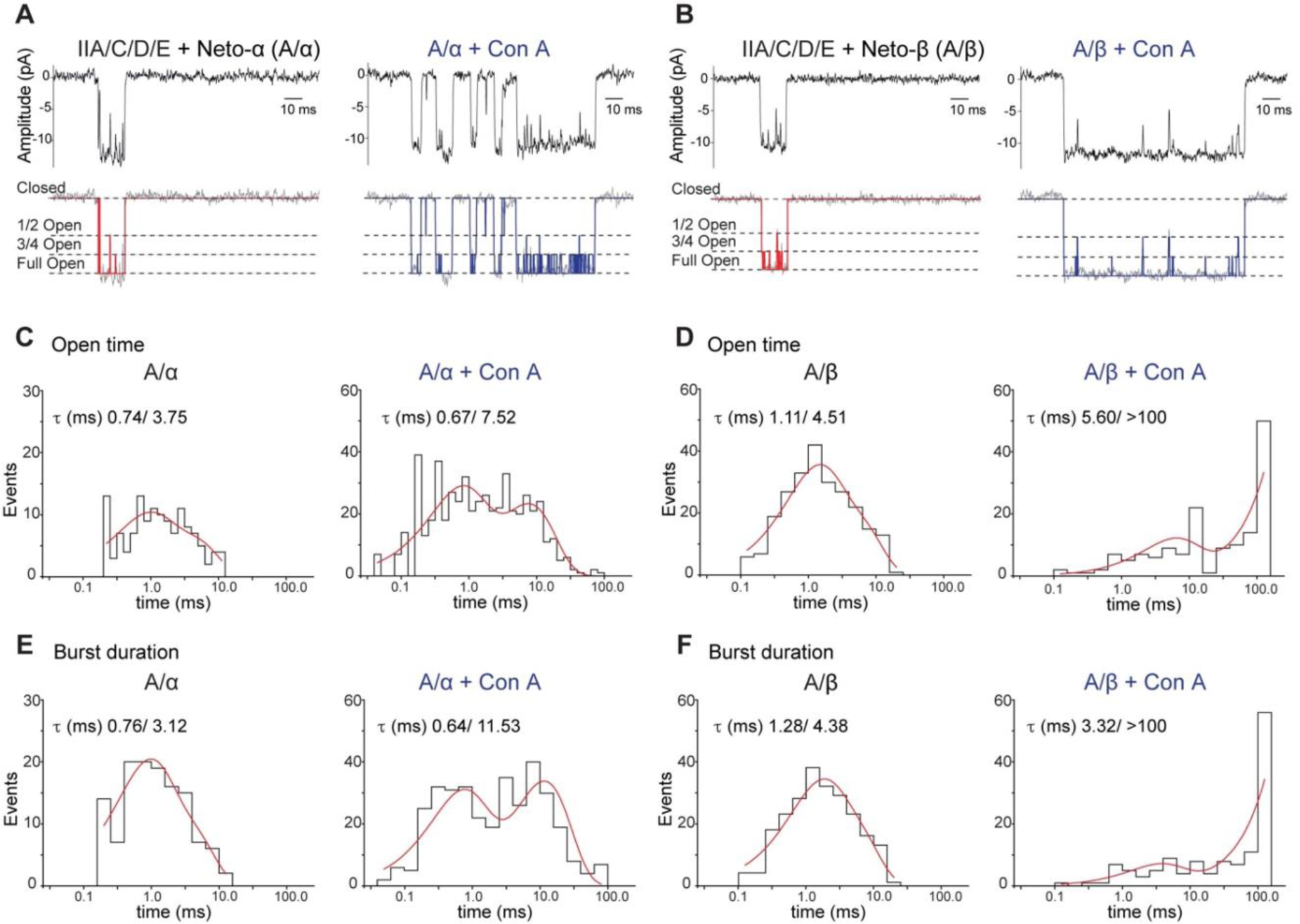
Analysis of single-channel kinetics for type-A receptors. (**A-B**) Filtered and idealized traces illustrating responses to 10 mM glutamate applied for 100 ms to outside-out patches from HEK293T cells transfected with A/α and A/β receptor complexes; left: control patches, right: patches obtained after treatment of HEK cells with 0.6 mg/ml Con A for 10 min; idealized traces, shown in red and blue, respectively, reveal the full open state and two subconductance states, 1/2 open and 3/4 open, as indicated by dotted lines. (**C-D**) open time histograms for A/α or A/β receptor complexes for control patches (left) and patches after treatment with Con A (right) fit with the sum of two exponential functions as indicated. (**E-F**) burst duration histograms fit with the sum of two exponential functions for A/α and A/β for control patches (left) and after treatment with Con A (right).

**Supplemental Table 1.** Analysis of single channel kinetics for recombinant type A/α and A/β receptor complexes for control patches, and patches obtained from HEK cells treated with Con A. Closed time histograms revealed bimodal distributions for all conditions, but except for A/α + Con A the number of events was too small to allow an accurate estimate of the lifetime. Log binned open time and burst length distributions were fit with the sum of two exponentials as shown below (Figure 3). For receptor complexes with Con A we observed single openings and bursts of openings which exceeded the length of the 100 ms application of glutamate. Examination of the lifetime of openings in a burst revealed longer openings for each subconductance state after application of Con A for both A/α and A/β (Supplemental figure S2).

| Recombinant iGluRs | Closed time (ms) | Open time (ms) | Burst length |
| --- | --- | --- | --- |
| A/α | Bimodal | 0.74, 3.75 | 0.76, 3.12 |
| A/α + Con A | 0.6, 3.8 | 0.67, 7.53 | 0.64, 11.53 |
| A/β | Bimodal | 1.11, 4.51 | 1.28, 4.38 |
| A/β + Con A | Bimodal | 5.60, > 100 * | 3.32, > 100 ** |

\* Openings of duration in excess of 100 ms were observed
\*\* Bursts of duration in excess of 100 ms were observed

**Supplemental Table 2.** Analysis of single channel kinetics reported in prior studies on native extrajunctional receptors expressed in *Drosophila* muscle. NR indicates Not Reported

| Native iGluRs | Closed time ms | Open time ms | Burst Length |
| --- | --- | --- | --- |
| Nishikawa 1995 | Tau crit 1 ms | NR | 0.069, 0.9, 4.9 |
| Broadie 1993 | NR | 0.23, 2.1 | NR |
| Heckmann 1995 | 0.073, 0.661 | 0.071, 1.81 | NR |

1. Nishikawa, Y. Kidokoro, Junctional and extrajunctional glutamate receptor channels in *Drosophila* embryos and larvae. *J Neurosci* **15**, 7905-7915 (1995). 2. K. S. Broadie, M. Bate, Development of the embryonic neuromuscular synapse of *Drosophila melanogaster*. *J Neurosci* **13**, 144-166 (1993). 3. M. Heckmann, J. Dudel, Recordings of glutamate-gated ion channels in outside-out patches from *Drosophila* larval muscle. *Neuroscience letters* **196**, 53-56 (1995).

**Supplemental figure S2.**
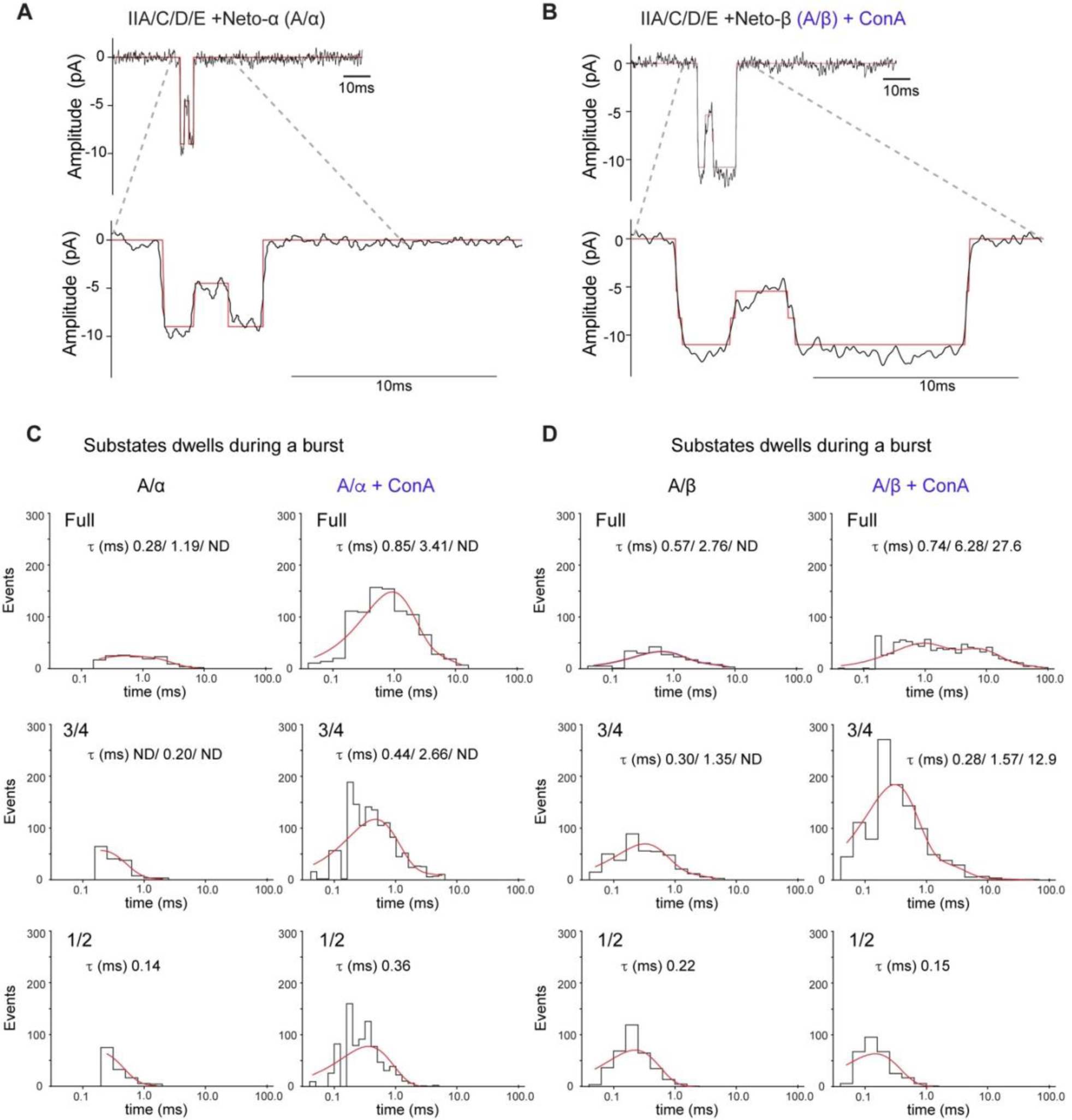
Modulation of substate activity by Con A. (**A-B**) Representative responses to 10 mM glutamate applied for 100 ms to outside-out patches from HEK cells transfected with A/α and A/β receptor complexes, as indicated. The idealized traces (in red) show transitions from the fully open state to the 1/2 level. (**C-D**) Dwell time histograms for substate occupancy during a burst for control patches (left panels), and for patches from HEK cells treated with Con A (right panels) for A/α (C) and A/β (D), fit with the sum of 1-3 exponentials of fast, intermediate and slow time constants, as indicated. ND-not detected.

**Supplemental figure S3.**
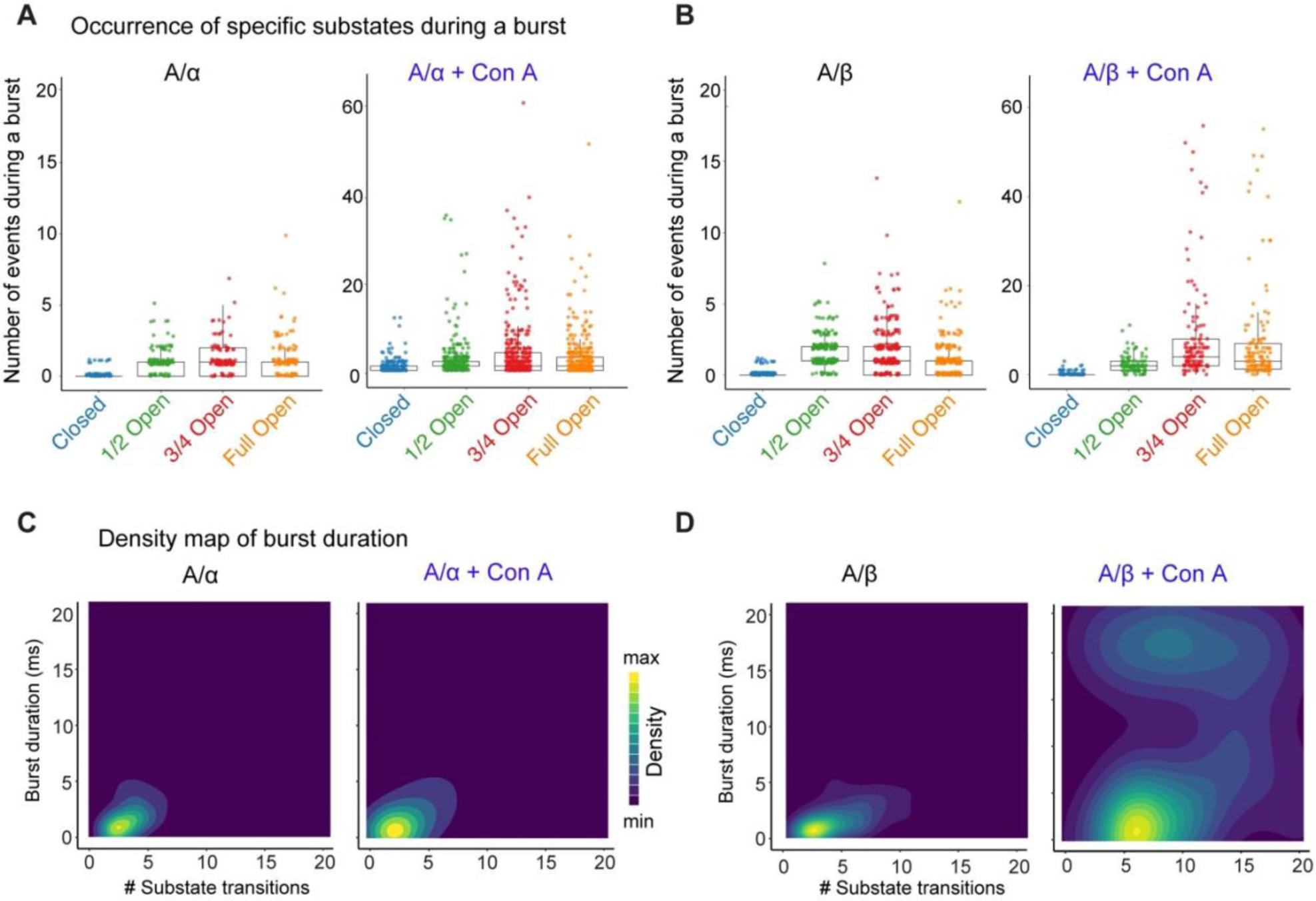
Distribution of substate transitions and burst characteristics. (**A-B**) Box plots indicating the number of specific events (substates occurrence) during a burst for A/α (**A**) and A/β (**B**) receptor channels before (left) and after (right) treatment with 0.6 mg/ml Con A for 10 min. The states closed, 1/2 open, 3/4 open, and fully open are color coded, as indicated. The boxes capture the interquartile range (IRQ) of each group’s distribution of values; horizontal lines denote median values; vertical lines extend to 1.5x IQR. (**C-D**) Density plots illustrating the relationship between burst duration and the number of transitions among substates for the A/α and A/β receptor complexes as indicated, in the absence (left) of presence (right) of Con A.

Analysis of the event list generated by the idealization process revealed subtle differences between channels containing Neto-α *versus* Neto-β (Supplemental figure S2). For example, A/α spends more time in the 1/2 open substate than A/β, which appears to spend a longer time in the 3/4 open substate. Application of Con A expanded these differences: A/α channels visit the closed and 1/2 open subconductance states very often, while A/β channels rarely transition into these states and instead spend most of the time in the 3/4 open and full open substates (Supplemental figures S2 and S3). The distribution of burst duration *versus* the number of transitions among substates further captured the different channel behaviors and indicated that A/α channels undergo ∼50-100 substate transitions within a burst (Supplemental figure S3C-D). Since all the substates are well represented during a burst, we conclude that, upon desensitization block, A/α channels transit between all subconductance states. In contrast, approximately 50% of A/β channel bursts showed few substate transitions, while the other 50% had ∼100 transitions per burst, suggesting modal gating. Due to brief lifetime of openings observed for type-B receptors (Figure 2B) we did not attempt a similar single channel kinetic analysis, but note that inspection of the raw data suggests that the burst length of B/α is shorter than for B/β similar to the behavior observed for A/α and A/β.

### Use dependent block by external philanthotoxin

Ca^2+^-permeable vertebrate kainate receptors are blocked by extracellular philanthotoxin (PhTx), a polyamine toxin derived from wasp venom (Eldefrawi et al. 1988; Bahring and Mayer 1998). Because native *Drosophila* NMJ iGluRs have high Ca^2+^-permeability (Chang et al. 1994) we tested the effects on PhTX on multi-channel outside-out patches at a concentration of 1 µM, while recording the response of type-A and type-B receptors to glutamate (Figures 4A and 4D). Before application of PhTx the mean charge transfer in response to 100 ms applications of 10 mM glutamate was: A/α, 4.08 ± 1.13 pC; A/β, 4.68 ± 1.77 pC; B/α, 0.17 ± 0.04 pC; B/β, 0.40 ± 0.14 pC. Similar to the macroscopic currents shown in Supplemental figure S1, we observed trial to trial amplitude variations for all channel combinations. Inhibition by PhTx developed slowly, τ_onset_ 78 s and 56 s for A/β and B/β, respectively (Supplemental figure S4), but at equilibrium, PhTx substantially reduced the charge transfer in response to a 100 ms application of glutamate to 1.6% ± 0.2 (n = 5) and 1.2% ± 0.3 (n = 5) of control, for type A/α and A/β channels respectively. For type-B channels, block at equilibrium was weaker, with the charge transfer reduced to only 16.7% ± 2.5 for B/α (n = 4) and 25.3% ± 0.9 for B/β (n = 4) compared to control. This difference in PhTx-induced block between type-A and type-B receptors resembles that observed in prior experiments using *Xenopus* oocytes to study block by the structurally related channel blocker argiotoxin (Han et al. 2015).

**Figure 4.**
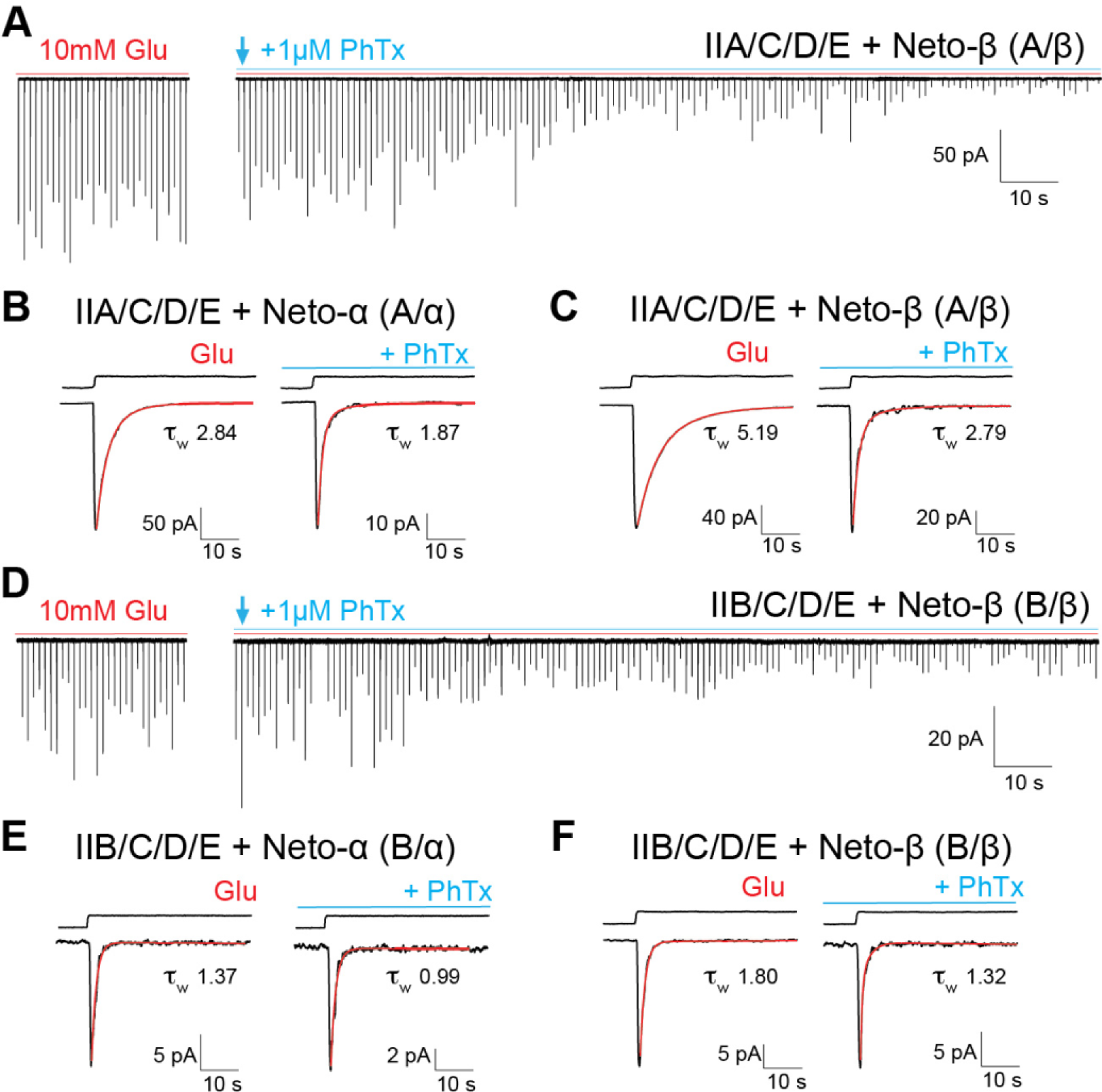
Slow onset of block by external PhTx. (**A** and **D**) Representative traces for type-A and type-B receptors showing responses recorded from outside-out patches to 10 mM glutamate applied for 100 ms at an interval of 1 s before (red line) and after the application of 1 µM PhTx (blue line). The amplitude variation is due to differences in the number of channels activated from trial to trial. (**B** and **C, E** and **F**) Averages of 20 responses before PhTx or starting 5 s after the onset of PhTx application for (**B**) A/α; (**C**) A/β; (**E**) B/α; (**F**) B/β. Red lines show fits of double exponential functions, and reveal faster decay in the presence of PhTx. The holding potential was -60 mV; back lines above the response to glutamate show open tip potentials.

**Supplemental figure S4.**
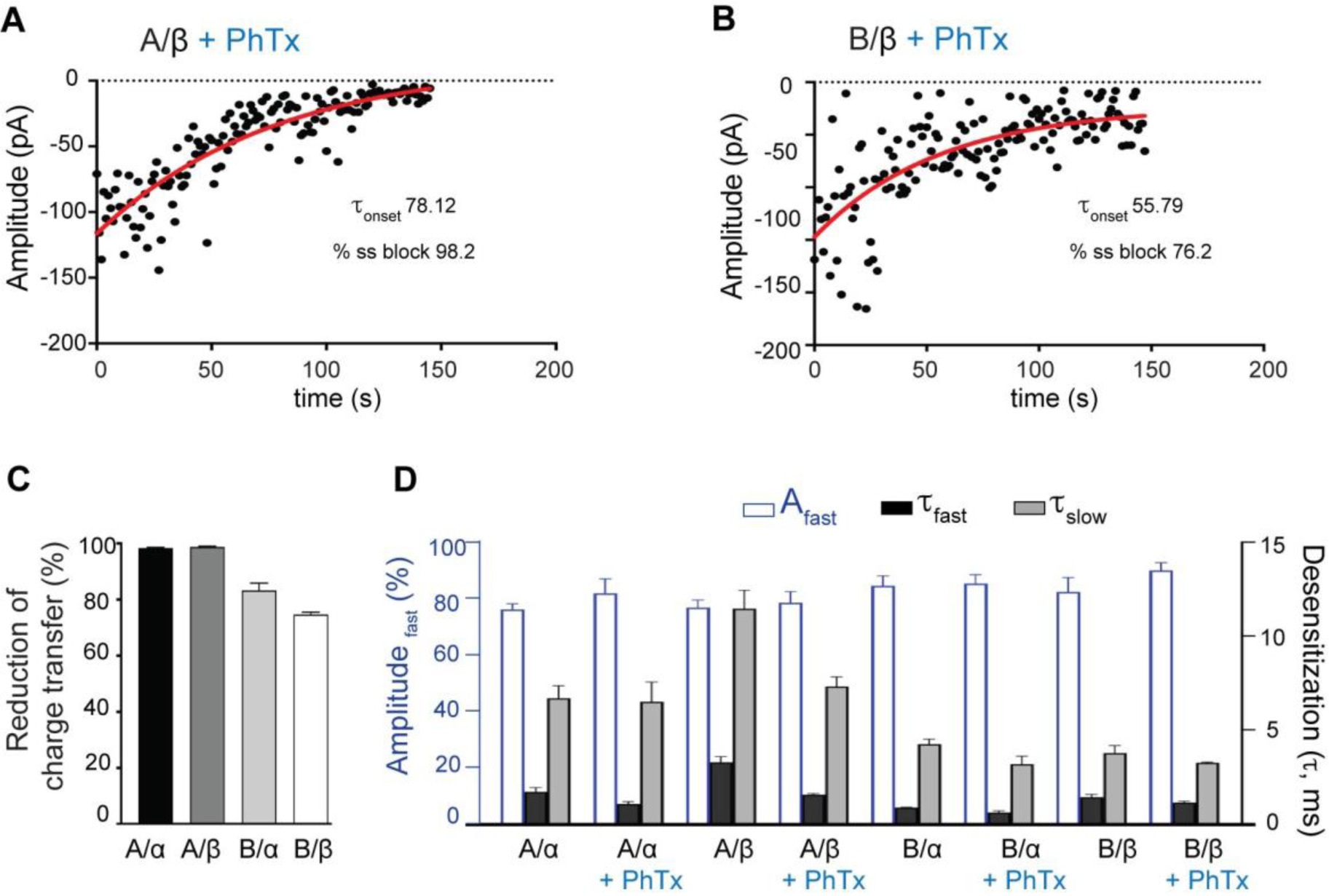
Slow onset of block by PhTx. (**A-B**) Data points indicate the amplitude of sequential responses to 10 mM glutamate applied for 100 ms at intervals of 1 s for A/β and B/β. The amplitude variation is due to differences in the number of channels activated from trial to trial. Red lines show single exponential fits to the decay of the response to glutamate due to onset of block by 1 µM PhTx. (**C**) The extent of block by PhTX at equilibrium, estimated from the change in charge transfer with respect to control, where a value of 100% indicates complete block, with values A/α, 98.4 ± 0.23 % (n = 5); A/β, 98.8 ± 0.30 % (n = 5); B/α, 83.31 ± 2.53 % (n = 5); B/β, 74.70 ± 0.86 % (n = 5). (**D**) Fits to the decay of the response to 100 ms applications of 10 mM glutamate (fit with the sum of two exponentials) before and after application of PhTx, showing mean values for %A_fast_, τ_fast_, and τ_slow_ for the indicated combinations of type-A and type-B complexes with Neto-α and Neto-β.

The reduction in charge transfer by PhTX results from two effects. First, the number of channels that open in response to glutamate progressively decreases in the presence of PhTx (Figures 4A and 4D). Second, the rate of decay of the response to glutamate, which in control conditions results from the onset of desensitization, increases; this is most likely due to a combination open channel block by PhTx combined with the onset of desensitization. Indeed in the presence of PhTx, the kinetics of decay were substantially faster, as estimated from the average response to 15-30 applications of glutamate recorded immediately before and after the start of the application of toxin, τ_w_ A/α 2.84 ± 0.20 ms before and 1.87 ± 0.08 ms after PhTx (n = 5); τ_w_ B/α 1.37 ± 0.14 ms *versus* 0.99 ± 0.09 (n = 4); τ_w_ A/β 5.19 ± 0.39 ms *versus* 2.79 ± 0.23 (n = 5); and τ_w_ B/β 1.80 ± 0.15 ms *versus* 1.32 ± 0.07 (n = 4). Most of this change was due to an increase in the fast component of decay of the response to glutamate (Table 2). The slow rate of onset of block by PhTx (Figures 4A and 4D) likely reflects open channel block and the low open channel probability (Supplemental figure S1) convolved with the limited time that the channel spends in the open state due to rapid onset of desensitization in response to glutamate.

**Table 2.**
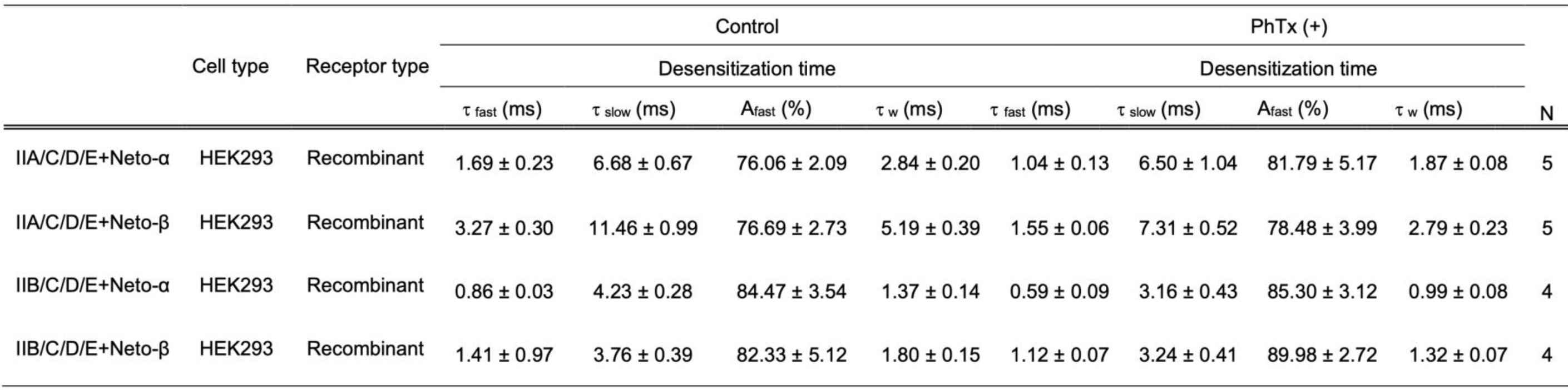
Decay kinetics for responses to 100 ms applications of 10 mM glutamate recorded from the same outside patch before and in the continuous presence of 1 µM PhTx. Responses were fit with the sum of two exponentials, revealing a 1.3 to 2.1-fold increase in the rate for the fast component of decay in the presence of PhTx.

### Native NMJ iGluRs

Differences between the properties of native and recombinant neurotransmitter receptors have been instrumental in the discovery of novel auxiliary proteins and synaptic modulators (Jackson and Nicoll 2011). *Drosophila* NMJ iGluR receptor channels are assembled from either of two alternative subunits, GluRIIA and GluRIIB, combined with two Neto isoforms. Null mutants are available for each of the four variants (DiAntonio et al. 1999; Kim et al. 2015; Han et al. 2020), permitting the *in vivo* isolation of individual receptor types with a single Neto isoform; this facilitates a direct comparison with recombinant iGluR/Neto channels expressed in HEK293T cells. We generated third instar larvae with single copies of either the *GluRIIA* or *GluRIIB* genes, and either *neto-α* or *neto-β*. Mutants with one or two copies of *GluRIIB* and *neto-α* (*neto-β^null^; GluRIIA^null^)* are embryonic lethal and thus could not be studied.

We recorded from outside-out patches obtained from the larval muscle membrane (muscle 6, abdominal segment 3) of animals with defined receptor complexes and compared the deactivation kinetics of responses to 1 ms applications of 10 mM glutamate with those of recombinant receptors of the same subunit composition expressed in HEK cells. We found that native, extrajunctional channels have slower deactivation kinetics than recombinant receptors (Figure 5A-B and Figure 1 and Table 1), A/α τ_off_, 1.57 ± 0.27 ms in muscle patches *versus* 0.64 ± 0.07 ms in HEK cells, and A/β τ_off_, 1.60 ± 0.29 ms *versus* 0.90 ± 0.11 ms, (n = 5 or 6). A less pronounced but similar trend was observed for B/β: recombinant receptors had faster deactivation τ_off_, 0.58 ± 0.06 ms, than the native extrajunctional complexes τ_off_, 0.97 ± 0.24 ms (Figure 5C). In contrast the single conductance γ: A/α, 169.3 ± 2.9 pS; A/β, 171.1 ± 3.51 pS; B/β, 172.6 ± 3.5 pS for native receptors was not different from values obtained for recombinant receptors. We next recorded mEJCs at the larval NMJ in mutants with defined receptor complexes and estimated the decay time for synaptic currents fit with single exponential functions (Figure 5D-F). The mEJC decay time constant was much slower than the deactivation time constant for both type-A and type-B extrajunctional receptors, τ_mEJC_: A/α, 4.20 ± 0.42 ms (n = 5); A/β, 6.50 ± 0.28 ms (n = 6); B/β, 4.92 ± 0.23 ms (n = 5). The slow decay cannot be explained by asynchronous release at multiple junctional sites since we recorded miniature EJCs. Instead, our data suggest that either the clearance of glutamate from the synaptic cleft is slow compared to other synapses, or that additional auxiliary membrane proteins and/or cytoplasmic modulatory proteins are present and impact the gating of synaptic receptors.

**Figure 5.**
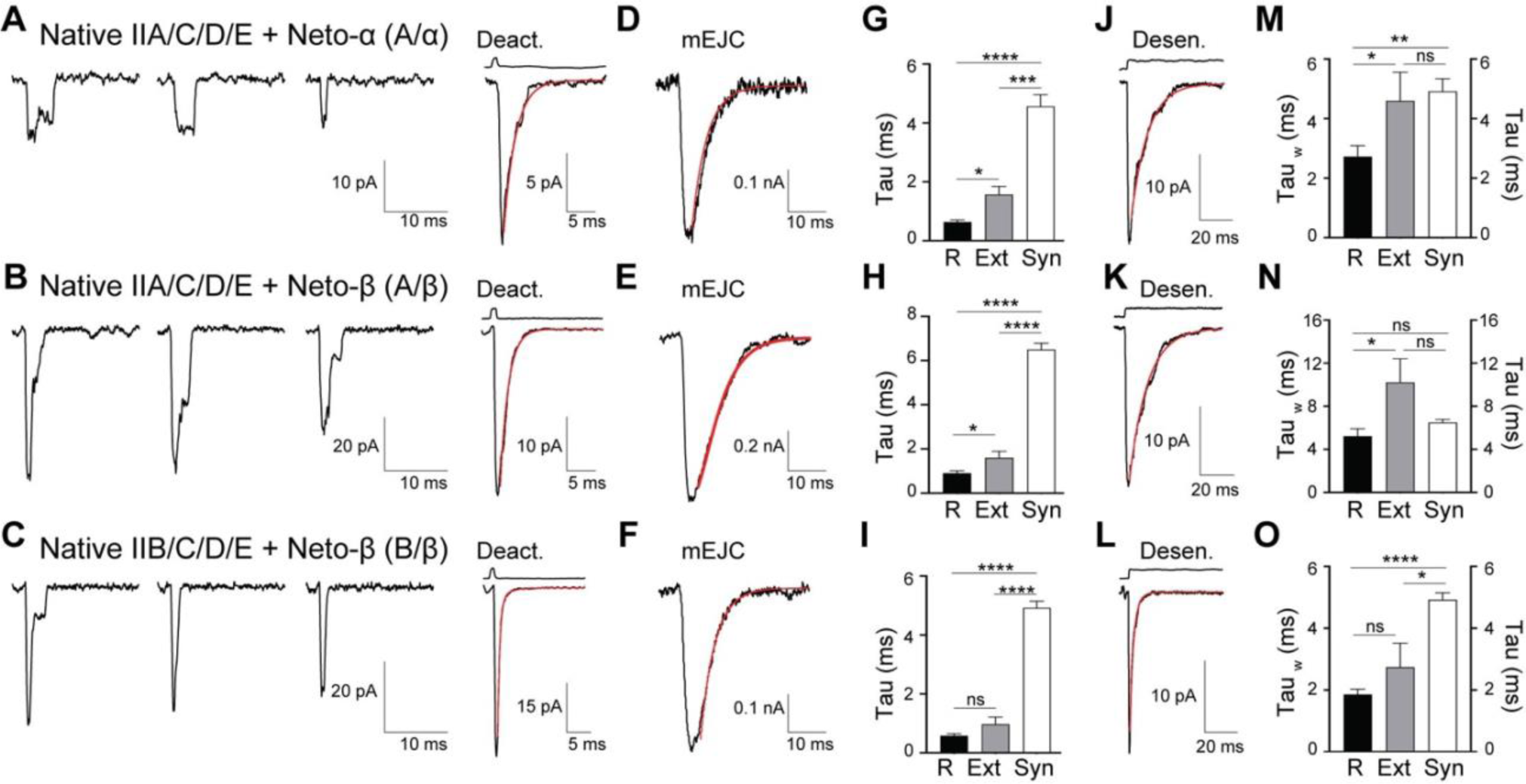
Distinct receptor properties observed for native and recombinant receptors. (**A-C**) Representative individual traces (left) and the average of 20-40 responses (right) to 10 mM glutamate applied for 1 ms to outside-out patches of native, extrajunctional receptors excised from body-wall muscles of third instar larvae with distinct iGluR/Neto complexes, as indicated. The red line shows time course of deactivation fit by an exponential function; open tip junction currents measured at the end of the experiments are shown at the top. (**D-F**) Miniature EJCs recorded from larvae of the same genotypes fit with single exponential functions. (**G-I**) Comparisons of deactivation kinetics for recombinant receptors expressed in HEK cells (R), native, extrajunctional receptors from larval muscle membranes (Ext) and mEJCs (Syn). (**J-L**) Average of 20-40 responses to 10 mM glutamate applied for 100 ms to outside-out patches of native, extrajunctional receptors for the same genotypes; the red line shows fits of the sum of two exponential functions; open tip junction currents measured at the end of the experiments are shown at the top. (**M-O**) Comparisons of desensitization kinetics for recombinant receptors expressed in HEK cells (R), native, extrajunctional receptors from larval muscle membranes (Ext) and mEJCs (Syn). Data are represented as mean ± SEM. **** p<0.0001, *** p<0.001, ** p<0.01, *p<0.05, ns, p>0.05.

To compare the kinetics of mEJCs with the kinetics of desensitization we next recorded responses to 100 ms applications of 10 mM glutamate from native extrajunctional receptors. This revealed that the desensitization time constant for A/α, 4.60 ± 0.96 ms (n = 5) was nearly identical to the mEJC decay time constant of 4.20 ms. The desensitization time constant for native A/β, 9.4 ± 2.6 ms (n = 6) was 1.4-fold slower than the mEJC decay time constant of 6.5 ms. Curiously, the desensitization time constant for native B/β, 2.74 ± 0.78 ms (n = 5) was 1.8-fold faster than the mEJC decay time constant of 4.9 ms.

## Discussion

The fly NMJ has been used extensively for genetic analysis of synapse development and homeostasis. Since the trafficking and synaptic stabilization of NMJ iGluRs depends on receptor activity (Marrus and DiAntonio 2004; Petzoldt et al. 2014), our lack of an understanding of how receptor properties are modulated has been a decades long gap in the field. Here we address this issue by measuring the function of recombinant *Drosophila* NMJ iGluRs expressed in HEK293T cells and compare their gating with native extrajunctional and synaptic receptors. We find that the kinetic properties of type-A and type-B receptors are distinct, with differential modulation by Neto isoforms adding further diversity to their kinetics. In addition, different from vertebrate kainate receptors, Neto is absolutely essential for the function of *Drosophila* NMJ iGluRs. Our data build on previous findings obtained for recombinant NMJ receptors expressed in *Xenopus* oocytes and further explain why Neto is essential *in vivo* for NMJ functionality and organism viability (Kim et al. 2012; Han et al. 2015).

### iGluR kinetics and synaptic transmission at the larval NMJ

Building on the pioneering discovery by Katz and his colleagues of quantal synaptic transmission (Fatt and Katz 1952), a central tenet of synaptic physiology is that the rate of closure of ion channels activated by neurotransmitters determines the time course of the synaptic response. This was first established for the vertebrate NMJ for which decay of the endplate current was shown to be determined by the intrinsic rate constant for closure of nicotinic acetylcholine receptors (Magleby and Stevens 1972). At vertebrate neocortical synapses, which similar to the *Drosophila* NMJ use glutamate as a neurotransmitter, the time constant for channel closure, τ_off_ 2.1 ms, is the same as the mEPSC decay time constant, τ_mEPSC_ 2.3 ms (Hestrin 1992), with slower mEPSC decay time constants in hippocampal neurons τ_mEPSC_ 4-8 ms (Hestrin et al. 1990). In our experiments we found that the deactivation kinetics for recombinant *Drosophila* NMJ iGluRs were very fast, with sub ms kinetics for both type-A and type-B receptor complexes with Neto-α, τ_off_ 0.64 and 0.42 ms, respectively. Slightly less rapid deactivation was recorded for type-A and type-B receptor complexes with Neto-β, τ_off_ 0.90 and 0.58 ms, respectively. For outside out patches obtained from the muscle of larvae with genetically controlled receptor composition, deactivation was 2-fold slower compared to recombinant iGluRs for both type-A and type-B receptor complexes, τ_off_ 1.57 ms for A/α, 1.60 ms for A/α, and 0.97 ms for B/β. Although these rates are faster than measured for AMPA receptor deactivation and glutamatergic mEPSC decay in hippocampal and cortical neurons, sub-millisecond deactivation and mEPSC decay have been recorded from neurons in brain stem auditory pathways and cerebellar Purkinje cells, likely reflecting differences in the subunit and auxiliary proteins present in different iGluR complexes (Barbour et al. 1994; Raman et al. 1994). The different kinetics of deactivation and desensitization for recombinant receptors and native extrajunctional channels observed in our experiments may reflect differences in post-translational modifications or lipid microenvironments surrounding these channels in HEK cells *versus* larval muscle. Alternatively, as yet unidentified additional accessory subunits or trans-synaptic proteins might modulate receptor function.

We found that the decay time constants of mEJCs, recorded from larvae with genetically controlled receptor and Neto splice variant composition, were 3-5 times slower than for deactivation measured for native extrajunctional receptor-Neto complexes of the same composition. By contrast, the rate of decay mEJC was nearly identical to the rate of desensitization for A/α, τ_mEJC_ 4.2 ms, τ_w_ 4.6 ms; for A/β the rate of desensitization was 1.4-fold slower than the decay of the synaptic current, τ_mEJC_ 6.5 ms, τ_w_ 9.4 ms; unexpectedly, for B/β the rate of desensitization was 1.8-fold faster than the synaptic current, τ_mEJC_ 4.9 ms and τ_w_ 2.7 ms. Because animals with B/α receptors die during late embryogenesis, and are unable to hatch, it is likely that B/α receptors deactivate and desensitize extremely rapidly and cannot sustain normal synaptic transmission and muscle contraction. The rates of desensitization of A/α and A/β complexes were significantly slower than for the previously reported A/α+β complexes, τ_w_ 4.6 ms and 9.4 ms *versus* 19 ms, respectively (DiAntonio et al. 1999); this may reflect an interaction between Neto-α and Neto-β when bound to the same cluster of heterotetrameric receptors or may be caused by the association of A/α+β complexes with other modulators. In the larval muscle, Neto-β is ∼10 fold more abundant than Neto-α, but both isoforms contribute to the proper assembly of the PSDs (Ramos et al. 2015; Han et al. 2020).

Overall, our results suggest that desensitization plays a major role in determining the kinetics of synaptic transmission at the larval NMJ. This result explains previous observations that the K661E mutation within the M3-S2 linker of GluRIIA, which effectively blocks desensitization of vertebrate homomeric GluA2 (Yelshansky et al. 2004), slows synaptic current decays at *GluRIIA^K661E^*-containing larval NMJs (Petzoldt et al. 2014). By contrast, the mEPSC decay time constant in *GluRIIA* mutants that correspond to fast-desensitizing vertebrate GluA2 mutants resembles the fast kinetics of type-B receptors. This critical role for desensitization in the kinetics of synaptic transmission may constitute an adaptation of larval NMJ to the high concentration of glutamate (1.8 - 2.0 mM) in the hemolymph, which is actively maintained (Augustin et al. 2007; He et al. 2023). Mutations that reduce the hemolymph glutamate concentration (to ∼1 mM) or culturing larvae in glutamate-depleted conditions trigger a 250-500% increase in the levels of postsynaptic type-A and B receptors. Interestingly, application of Con A to glutamate-depleted larvae completely blocked the increase in iGluR clustering (Augustin et al. 2007), indicating that glutamate-mediated suppression of iGluR synaptic recruitment might depend on receptor desensitization. Of note, alterations in ambient extracellular glutamate also dramatically alters glutamatergic neurotransmission in cultured vertebrate hippocampal neurons (Lissin et al. 1999). We are not aware of any reports for which the synaptic current decay at vertebrate synapses is determined exclusively by the rate of desensitization. However, for some vertebrate CNS glutamatergic synapses desensitization has been shown to contribute to a slow component of synaptic currents (Barbour et al. 1994; Koike-Tani et al. 2005). In addition, desensitization shapes the response to multiquantal neurotransmitter release, and to paired pulse stimulation at vertebrate glutamatergic synapses (Trussell et al. 1993).

In our experiments, mEJCs were recorded using two-electrode voltage clamp which revealed rise times of around 1.5 ms and decay time constants of 4-6 ms, which varied with subunit composition. Previous recordings from control animals, which express both type-A and type-B receptors or *GluRIIA^null^* mutants, with only type-B receptors, reported decay time constants of 7.73 ± 1.52 ms and 2.95 ± 0.65 ms, respectively (Petzoldt et al. 2014). Furthermore, a survey of the literature, for which we manually digitized published records for mEJC decay obtained using extracellular focal recording (Stewart et al. 1994; Cooper et al. 1995; Heckmann and Dudel 1998; Dawson-Scully et al. 2007; Karunanithi et al. 2018), yielded a mean value for τ_mEJC_ of 4.37 ± 0.17 ms (n = 7), range 2.5 to 7.7 ms, similar to values recorded in the present experiments. In prior studies the rise time of mEJCs measured using focal recording varied from 0.35 to 1.1 ms (Heckmann and Dudel 1998; Paul et al. 2015) slightly faster than we and others (Petzoldt et al. 2014) recorded using two-electrode voltage clamp. Taken together, our results suggest that the time course of decay of the concentration of glutamate in the synaptic cleft of the larval NMJ is slow compared to conventional synapses and determined by desensitization and not deactivation.

### Modulation of desensitization by Con A and single channel kinetics

Similar to vertebrate kainate receptors (Huettner 1990; Wong and Mayer 1993), application of Con A consistently attenuated (type A/α) or blocked (A/β, B/α and B/β) desensitization, and increased the ratio of charge transfer, open time and burst duration of single channels expressed in HEK cells. The initial characterization of *Drosophila* NMJ iGluRs expressed in *Xenopus* oocytes was only possible after application of Con A (Han et al. 2015), and it remained possible that receptors expressed in the absence of Neto were activated by glutamate, but desensitized too rapidly to detect using oocyte recording. In the experiments reported here we used outside out patches with sub ms solution exchange and found that in the absence of Neto, *Drosophila* NMJ iGluRs are not activated by glutamate on a physiological time scale; this explains why *neto^null^* mutants are embryonic lethal (Kim et al. 2012). We cannot exclude the possibility that receptors which traffic to the membrane desensitize so rapidly in the absence of Neto that the channel does not open, as found for the AMPA receptor S750D and E755A mutants (Partin et al. 1996; Horning and Mayer 2004).

Prior studies on native *Drosophila* NMJ iGluRs have largely focused on measurements of single channel conductance (Broadie and Bate 1993a; Nishikawa and Kidokoro 1995; Heckmann and Dudel 1997; DiAntonio et al. 1999). There is good agreement among these studies with results obtained in the present experiments, which reveal a single channel conductance of 160 to 170 pS, with no difference between native and recombinant type-A and type-B receptors or Neto isoforms. Our experiments on recombinant *Drosophila* NMJ iGluRs revealed frequent transitions to subconductance states of 75% and 50% of the main state when activated by 10 mM glutamate. This was observed for both control patches and for patches from cells pretreated with Con A, but were easier to detect in the latter condition due to an increase in the open time. The presence of subconductance states is a widely observed feature of the gating of vertebrate AMPA and kainate receptors (Rosenmund et al. 1998; Daniels et al. 2013; Coombs and Cull-Candy 2021; Baranovic et al. 2022) and it is surprising that this has not been reported before for native extrajunctional *Drosophila* NMJ iGluRs. Inspection of published raw data from prior single channel recording experiments reveals hints of substate activity, though it is possible that this results from brief transitions between fully open and closed states that were not resolved (Heckmann and Dudel 1995; Nishikawa and Kidokoro 1995; Heckmann and Dudel 1997; DiAntonio et al. 1999). In the future, it will be interesting to expand the analyses of substates and compare the behavior of recombinant and extrajunctional receptors at physiological glutamate concentrations.

For measurement of single channel lifetimes we used a cutoff value of 200 µs for generating the list of idealized events, and a critical time of 2 ms for burst length analysis; as a consequence we did not resolve brief events with µs lifetimes reported in prior studies on native receptors (Heckmann and Dudel 1995; Chang and Kidokoro 1996). Due to the limited number of single channel events recorded in control patches as a result of rapid desensitization, combined with their short duration, we did not calculate closed time distributions, and we limited our analysis to recombinant type-A receptors for which openings were well resolved. For control patches the open time distributions were comparable to those reported for wild type extrajunctional receptors, as summarized in Supplemental table 1 (Broadie and Bate 1993b; Heckmann and Dudel 1995; Nishikawa and Kidokoro 1995), while for patches from HEK cells treated with Con A there was a substantial increase in open time. For the burst length distribution there is only a single observation in the literature for native iGluRs, with medium and long duration events of life time 0.9 and 4.9 ms (Nishikawa and Kidokoro 1995)(Supplemental table 2); these values are comparable to those obtained for recombinant receptors A/α, 0.76 and 3.12 ms and A/β, 1.28 and 4.38 ms (Table 1). Treatment with Con A increased the lifetime of long duration openings for A/α nearly 4-fold; for A/β the lifetime of long duration events in the open time and burst duration distributions exceeded 100 ms, resulting in tail currents following removal of glutamate (Figures 2C, 3C and 3E, and Table 1). In addition we analyzed the open time distribution of substates within bursts and found additional subtype specific differences (Figure 3 - figures supplement 1 and 2, and Table 1).

### Channel block by polyamine toxins

Similar to vertebrate Ca^2+^-permeable receptors (Bowie and Mayer 1995; Kamboj et al. 1995) we previously found that recombinant *Drosophila* NMJ iGluRs expressed in *Xenopus* oocytes show biphasic rectification and were blocked by Argiotoxin (Han et al. 2015). Prior studies on *Drosophila* NMJ synaptic responses also revealed biphasic rectification (Broadie and Bate 1993b; Nishikawa and Kidokoro 1995), with similar voltage dependence to that produced by polyamines indicating that channel block by cytoplasmic polyamines modulates synaptic transmission *in vivo*. This behavior is likely critical to flies, which live on rotten fruits and tolerate large amounts of polyamines in their food. Interestingly, disruption of the polyamine metabolism in the fly muscles causes progressive locomotor defects that can be rescued by polyamine supplementation (Coni et al. 2021).

Block of *Drosophila* NMJ iGluRs by philanthotoxin has particular relevance for studies of synaptic plasticity at the NMJ in response to application of PhTX. In our experiments we found that PhTx produces a slow onset cumulative block of responses to sequential applications of glutamate, as well as an acceleration in the rate of decay of individual responses to glutamate (Fig. 5 and Table S2). At the *Drosophila* NMJ, manipulations that reduce postsynaptic iGluR activity trigger a compensatory increase in neurotransmitter release (DiAntonio et al. 1999; Frank et al. 2006). This form of plasticity has been intensely studied using two experimental paradigms: (1) a chronic (developmental) response in *GluRIIA^null^* mutants, and (2) the acute response to PhTx (20 μM) applied to dissected larval fillets. The two settings appear to elicit genetically distinct responses (Frank 2014). Our findings that external application of PhTx blocks both type-A (>98%) but also type-B receptors (by ∼80%) indicate that both type-A and type-B receptors are impaired during acute potentiation by PhTx application. Thus the two different plasticity paradigms differ in the time frame, developmental *versus* acute, but also in the nature of receptors affected, more specifically type-A receptors are absent in the developmental paradigm, while both type-A and type-B mediate the acute response. Since type-A and type-B receptors are recruited and stabilized at the NMJ via genetically distinct mechanisms (Parnas et al. 2001; Liebl and Featherstone 2008; Petzoldt et al. 2014; Ramos et al. 2015; Sulkowski et al. 2016) disruption of their activity is expected to elicit different compensatory mechanisms.

### A central role for Neto at the *Drosophila* NMJ

To build a functional NMJ, iGluRs must assemble and traffic to the cell surface, and then be recruited to and stabilized at synaptic sites. *Drosophila* Neto controls the synaptic recruitment and composition of iGluRs at the larval NMJ (Kim et al. 2012; Kim et al. 2015; Ramos et al. 2015; Sulkowski et al. 2016). Specifically, Neto engages in extracellular interactions that cluster postsynaptic iGluRs in large aggregates (Kim et al. 2015); the intracellular domains of Neto sculpt the PSD size and type-A *versus* type-B receptor composition (Ramos et al. 2015; Han et al. 2020). *Drosophila* Neto is also critical for the function of NMJ iGluRs as revealed by our studies on recombinant receptors: In the absence of Neto, iGluRs are trafficked to the cell surface in both *Xenopus* oocytes (Han et al. 2015) and HEK cells, but fail to respond to glutamate; by contrast, co-expression with Neto permitted detection of rapidly desensitizing glutamate evoked responses (Figures 1 and 2). This requirement for Neto appears to be specific to heterotetrameric NMJ iGluRs; by contrast, similar to vertebrate KARs (Herb et al. 1992), the *Drosophila* KaiR1D subunit, which contributes to synapses in the fly brain, forms functional glutamate-gated homotetrameric channels independently of Neto (Li et al. 2016). Our results indicate that Neto-β and a minimal Neto, with no intracellular domain (Neto-ΔCTD), confer similar rates of deactivation and desensitization to type-A receptors (Figure 1), while the rate of desensitization of type-A receptors expressed with Neto-α is circa 2-fold faster. This suggests that the large intracellular domain of Neto-β functions primarily as an organizing scaffold at the NMJ, as previously proposed (Ramos et al. 2015), whereas Neto-α directly modulates gating. These findings are reminiscent of vertebrate Neto1 and -2, which differentially modulate iGluR gating properties, expanding their functional repertoire (Tomita and Castillo 2012).

In flies as in vertebrates, the iGluR (/Neto) complexes are further modulated by post-translational modifications that impact the synaptic distribution and the biology of these channels (Davis et al. 1998; Morimoto et al. 2009; Perry et al. 2022) Whether and how such modifications impact NMJ iGluR channel properties remains to be determined. Unlike vertebrate receptors, fly NMJ iGluRs have relatively short C-terminal tails, suggesting that Neto has additional roles in receiving and transducing signals concerning synapse activity and the network status. Also, postsynaptic iGluR/Neto complexes align likely via trans-synaptic columns with the presynaptic active zone scaffold, Bruchpilot (Muttathukunnel et al. 2022) and with presynaptic BMP signaling complexes (Sulkowski et al. 2016; Vicidomini and Serpe 2022); through such interactions Neto can relay postsynaptic signals to presynaptic effectors that mediate presynaptic responses.

The fly NMJ has been a powerful genetic model to investigate conserved mechanisms for synapse assembly, development and homeostasis. Disruption of kainate receptor biology at this synapse triggers a wide range of morphological and behavioral phenotypes, from a complete loss of postsynaptic receptors and synaptic structures, which causes embryonic paralysis and developmental lethality, to defects in the assembly and maintenance of PSDs, which generally induce locomotor deficits, and finally to subtle changes in postsynaptic composition and impairments in synaptic plasticity. Within this wide range of biological outcomes, the investigations on fly NMJ iGluRs and their modulation by Neto have the potential to reveal new functions for iGluRs and Neto and new modalities of regulation. Our study paves the way to parse out and elucidate the multiple functions and regulation of kainate receptors and their auxiliary proteins Neto.

## Materials and Methods

### Molecular constructs and expression of iGluRs in HEK293T cells

iGluR coding sequences placed between the Tolloid-related signal peptide (Serpe and O’Connor 2006) and an RGSH_6_ C-terminal tag, as previously described (Han et al. 2015), were subcloned into a cytomegalovirus expression vector, pRK5-IRES (Li et al. 2016). The pRK5-Neto constructs included full length Neto-α and Neto-β, derived from GH11189 and RE42119, respectively (Han et al. 2015), Neto-ΔCTD (M1-D478) (Ramos et al. 2015), and PM-Neto-ΔCTD (R^123^I and R^126^I) (Kim et al. 2015). HEK293T cells were cultured in Dulbecco’s Modified Eagle Medium (DMEM; Gibco) with 10% fetal bovine serum, 1% Glutamax and 1% Penicillin Streptomycin solution at 37°C in a 95% oxygen and 5% carbon dioxide incubator. HEK293T cells (2 x 10^5^/ml) were plated on 5 mm diameter coverglass coated with bovine collagen (Nutragen), attached for 24h, then transfected with pRK5-based constructs of iGluR subunits and Neto variants (1 µg total DNA/ ml cells) using ViaFect transfection reagent (Promega). The cells were incubated at 37°C for 3 hours, and then the temperature was decreased to 30°C. Outside-out patch-clamp recordings were performed 2-3 days after transfection at room temperature.

### Electrophysiological techniques

Outside-out patch recordings from HEK293T cells and muscle 6 (segment A3), with fast solution exchange achieved using four-bore glass tubing mounted on a P245.30 piezoelectric stack driven by a P-270 HVA amplifier (Physik Instrumente), were performed at room temperature as previously described (Horning and Mayer 2004). Recordings were made with thin-wall borosilicate glass pipettes (resistance, 3-6 MΩ). The external solution contained (in mM) 145 NaCl, 5.4 KCl, 1.8 CaCl_2_, 1 MgCl_2_, 5 HEPES (pH 7.3, osmolarity, 295 mOsm), to which 10 mM L-glutamate was added. The internal solution contained (in mM) 110 CsCl, 10 CsF, 0.5 CaCl_2_, 1 MgCl_2_, 10 HEPES, 5 CsBAPTA and 20 Na_2_ATP (pH 7.3). Open tip junction potentials recorded at the end of experiments typically had 10-90% rise times of < 300 µs, and data from patches for which responses to 10 mM glutamate had rise times > 400 µs were discarded. Concanavalin A (type IV), glutamate and Philanthotoxin-343 were purchased from Sigma-Aldrich.

### Fly strains and *in vivo* recordings

Four different types of mutations were used to study the gating properties of *Drosophila* NMJ glutamate receptor channels by receptor subunit and Neto isoform composition: *GluRIIA^SP16^*(Petersen et al. 1997), *GluRIIB[Mi03631*] (BDSC# 37066), *neto-α^null^*(Han et al. 2020) and *neto-β^null^* (Ramos et al. 2015). These *GluR* mutants were crossed with a deficiency covering both *GluRIIA* and *GluRIIB* loci, *Df(2L)cl^h4^ (Petersen et al. 1997)*, similarly *neto* alleles were crossed with a *neto^null^* mutant (Kim et al. 2012). To study the gating properties of extrasynaptic receptor channels using outside-out patch recording as described above, wandering third instar larvae were dissected in ice-cold, calcium-free hemolymph-like HL-3 saline, then incubated with 30 μg/ml collagenase type IV (Sigma-Aldrich) for 10 min, washed with calcium-free HL-3 saline and moved to the recording chamber. The calcium-free HL-3 saline contained (in mM): 70 NaCl, 5 KCl, 20 MgCl_2_, 10 HCO_3_, 5 trehalose, 115 sucrose, 5 HEPES (pH 7.2). To examine the properties of synaptic receptor channels, two-electrode voltage-clamp recordings were performed on from muscle 6, segment A3 at room temperature as described previously (Qin et al. 2005). The recording solution was HL-3 with 0.5 mM CaCl_2_. Intracellular electrodes (borosilicate glass capillaries of 1 mm diameter) were filled with 3 M KCl with resistances ranging from 12 to 25 MΩ. Recordings were done from muscle cells with an initial membrane potential between –50 and – 70 mV, and input resistances of ≥ 4 MΩ. To record spontaneous miniature excitatory junction currents (mEJCs) the muscle cells were clamped to –80 mV. To calculate mEJCs mean amplitudes, 50–100 events from each muscle were measured and averaged using the Mini Analysis program (Synaptosoft).

### Data analysis

To calculate deactivation and desensitization time constants, 50 ∼ 100 representative responses were averaged and fit using a first order exponential function for deactivation

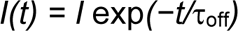

and a double exponential function for desensitization

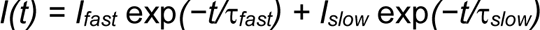

where *I_x_* is the peak current amplitude and τ_x_ is the corresponding decay time constant. To allow for comparison of decay times with published values fit with single exponential functions, weighted time constants τ_w_ were calculated using the formula

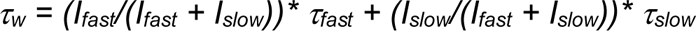

Current voltage plots recorded in the presence of spermine were fit with the sum of two Boltzman functions of the form, *G = G_max_ /[1+ exp((V_m_-V_b_)/k_b_)]+ G_max_ /[1+ exp((V_m_-V_p_)/k_p_)]*, where G_max_ is the conductance in the absence of block; G the conductance at membrane potential V_m_; V_b_ is the potential for half block; k_b_ the slope factor for block; V_p_ is the potential for half relief of block; k_p_ the slope factor for relief from block.

Single channel activity from outside-out patches was recorded using a gap-free protocol with an Axopatch 200B amplifier (Axon Instruments), digitized with a Digidata 1550 (Molecular Devices), sampled at 20 kHz, low pass filtered at 5 kHz, and collected using pClamp10.7 (Molecular Devices). Synaptic currents were recorded using a gap-free protocol with an Axoclamp 2B amplifier (Axon Instruments), digitized with a Digidata 1440A (Molecular Devices), sampled at 20 kHz, low pass filtered at 1 kHz, and collected using pClamp10.7 (Molecular Devices).

### Single-channel analysis

Single-channel recordings collected using pClamp10.7 were first exported in MATLAB format. Custom MATLAB scripts were implemented to (1) rename different experimental variables for compatibility with the idealization software, and to (2) extract 150 ms intervals from the recordings, including 25 ms before and 25 ms after the 100 ms glutamate application. The idealization process was performed using a PYTHON-based open source, single-channel analysis application, ASCAM (Baranovic et al. 2022). This included (1) baseline correction using an offset method within the intervals [0, 0.025] and [0.125, 0.150] seconds, (2) Gaussian filtering (2 kHz cutoff), followed by (3) idealization itself using the “Analysis > Idealize” function in ASCAM, with a multiple threshold crossing algorithm. Short-duration events were excluded by choosing a resolution of 200 μs. We first identified the fully open state for all single channels, then isolated two additional open substates by eye, which corresponded to ½ and ¾ of the open state. We never observed ¼ open substates. The table of idealized events was imported into pClamp, and the distribution of log binned dwell times fit with exponential functions by maximum likelihood using pClamp10.7. A custom R pipeline (available upon request) was used to evaluate of burst duration and open dwells for different subconductance states and the transitions between different substates.

## Acknowledgments

T.H.H, R.V., C.I.R., and M.S. were supported by Intramural Program of the NICHD, grants ZIA HD008914 and ZIA HD008869 awarded to M.S. We thank Dr. A. Plested for advice on use of ASCAM.

